# Serological screening in wild ruminants in Germany, 2021/22: No evidence of SARS-CoV-2, bluetongue virus or pestivirus spread but high seroprevalences against Schmallenberg virus

**DOI:** 10.1101/2022.02.21.481262

**Authors:** Kerstin Wernike, Luisa Fischer, Mark Holsteg, Andrea Aebischer, Anja Petrov, Katharina Marquart, Ulrich Schotte, Jacob Schön, Donata Hoffmann, Silke Hechinger, Antonie Neubauer-Juric, Julia Blicke, Thomas C. Mettenleiter, Martin Beer

## Abstract

Wildlife animals may be susceptible for multiple infectious agents of public health or veterinary relevance, thereby potentially forming a reservoir that bears the constant risk of re-introduction into the human or livestock population. Here, we serologically investigated 493 wild ruminant samples collected in the 2021/22 hunting season in Germany for the presence of antibodies against the severe acute respiratory coronavirus 2 (SARS-CoV-2) and four viruses pathogenic for domestic ruminants, namely the orthobunyavirus Schmallenberg virus (SBV), the reovirus bluetongue virus (BTV) and ruminant pestiviruses like bovine viral diarrhoea virus or border disease virus. The animal species comprised fallow deer, red deer, roe deer, mouflon and wisent. For coronavirus serology, additional 307 fallow, roe and red deer samples collected between 2017 and 2020 at three military training areas were included. While antibodies against SBV could be detected in about 13.6% of the samples collected in 2021/22, only one fallow deer of unknown age tested positive for anti-BTV antibodies and all samples reacted negative for antibodies against ruminant pestiviruses. In an ELISA based on the receptor-binding domain (RBD) of SARS-CoV-2, 25 out of 493 (5.1%) samples collected in autumn and winter 2021/22 scored positive. This sero-reactivity could not be confirmed by the highly specific virus neutralization test, occurred also in 2017, 2018 and 2019, i.e. prior to the human SARS-CoV-2 pandemic, and was likewise observed against the RBD of the related SARS-CoV-1. Therefore, the SARS-CoV-2-seroreactivity was most likely induced by another, hitherto unknown deer virus belonging to the subgenus *Sarbecovirus* of betacoronaviruses.

## Introduction

Wild ruminants, either free-ranging or raised in enclosures, can be infected by a wide range of infectious agents that are pathogenic for livestock animals or humans [1]. In case of livestock diseases, spillover from domestic ruminants to wildlife is commonly assumed to be the initial source of infection and, conversely, wild animals may subsequently develop into a reservoir bearing the risk for re-introduction of the disease into the livestock population.

In central Europe, the most common wild cervid species include the European roe deer (*Capreolus capreolus*), fallow deer (*Dama dama*) and red deer (*Cervus elaphus*), with roe deer being most closely related to the North American white-tailed deer (both subfamily *Capreolinae*) [2]. Besides, European bison (*Bison bonasus*) and the ovine species mouflon (*Ovis gmelini*) are abundant. All those species were shown to be susceptible for the reovirus bluetongue virus (BTV) and the orthobunyavirus Schmallenberg virus (SBV) [3–8], two arboviruses of major veterinary relevance that have in common the insect vector species responsible for virus transmission and the affected host animals. Both, BTV and SBV, are transmitted by *Culicoides* biting midges and predominantly infect ruminants [9,10]. In domestic ruminants, SBV may induce fever, diarrhoea or decreased milk yield in non-pregnant animals and abortion, stillbirth or the delivery of severely malformed offspring when naïve dams are infected during gestation [10]. BTV-infections are often inapparent or subclinical, but can also lead to a systemic haemorrhagic fever that results from vascular injuries affecting multiple tissues and organs and that is inducing a high mortality rate [11]. SBV was detected for the first time in late 2011 in the German-Dutch border region [12], and thereafter spread very rapidly through the European ruminant population [13]. By now, it is established in an enzootic status in Central Europe including Germany with patterns of wave-like virus re-circulation to a larger extent every two to three years [14,15]. In contrast, Germany was officially recognized free from BTV between 2012 and 2018. Prior to 2012, more precisely between 2006 and 2009, a large outbreak of BTV serotype 8 occurred in the domestic ruminant population, which was eventually controlled by mandatory vaccination [16,17]. New BTV cases have been recorded since December 2018 up to February 2021 [18]. In France, BTV-8 re-emerged already in 2015 and is circulating since then [19,20], representing a constant risk for virus spread into neighbouring regions or countries. In affected regions, besides major domestic ruminant species also wildlife could contribute to virus maintenance through their susceptibility to BTV infection, their high population density and through vector maintenance. Furthermore, in recent studies wild ruminants were shown to be excellent indicators for BTV circulation in a given area [21].

Apart from vector-borne viruses, wild and domestic ruminants also share susceptibility to a number of infectious diseases transmitted by direct contact. Depending on the dynamics of interaction and the particular pathogen, wild ruminants may maintain a pathogen independent of domestic animal populations through a sylvatic cycle, from which the pathogen in question might be transferred back to farmed animals. In case of the ruminant pestiviruses bovine viral diarrhoea virus (BVDV, syn. *Pestivirus A* and *B*) and border disease virus (BDV, syn. *Pestivirus D*), virus maintenance and transmission in domestic ruminants are predominantly driven by *in utero* infected, immunotolerant, persistently infected (PI) offspring. These PI animals shed high amounts of infectious virus throughout their life, as they are unable to mount a specific immune response against the virus strain they are infected with [22–24]. For BVDV, an eradication program is in place in Germany since 2011, which led to a considerable decrease in the prevalence of PI animals in the cattle population [25]. However, BVDV PI animals have been identified in a wide range of other mammals including cervid species [26,27] and, in Germany, anti-BVDV antibodies were detected in about 7.7 % of free-ranging deer in the 1990s [28], i.e. before the start of the mandatory nationwide control program. Therefore, there are concerns of potential pestivirus circulation in wild ruminants, that could lead to re-introduction into the domestic ruminant population.

From a public health perspective, wild ruminants are considered to be reservoir or maintenance hosts of multiple viral, bacterial, fungal or parasitic diseases [1]. Only recently, when different variants of concern of the betacoronavirus severe acute respiratory syndrome coronavirus 2 (SARS-CoV-2) or specific antibodies were detected in white-tailed deer [29–32], fears arose that cervid species could form an animal reservoir also for this virus. SARS-CoV-2, which belongs to the subgenus *Sarbecovirus* of the betacoronaviruses together with SARS-CoV-1 [33], is the causative agent of the COVID-19 pandemic that unfolded since the beginning of 2020 and led to millions of human deaths globally. Clinical signs in infected individuals range from asymptomatic infection to severe pneumonia-like, fatal disease. Since the beginning of the pandemic, the role of livestock and wildlife species as potential reservoir hosts was discussed. Natural SARS-CoV-2 infections linked to human exposure have been reported in American mink, ferrets, felines, canines and primates [34]. For domestic ruminants such as cattle, goat and sheep a very low susceptibly was demonstrated during experimental infection studies [35–37], and natural infections of cattle also seem to be a rare event [38]. However, SARS-CoV-2 or specific antibodies were detected frequently in free-ranging white-tailed deer (*Odocoileus virginianus*) in North America [29–31], but it remains unclear how these deer acquired the virus from humans. Under experimental conditions, white-tailed deer can be readily infected with SARS-CoV-2 and transmit the virus to conspecifics and, in case of pregnant dams, to the foetus [39,40]. Furthermore, the high seroprevalence observed in multiple US states and counties [30] suggests efficient natural intra-species transmission as well. *In silico* modelling suggested that additional deer species may be likewise susceptible to SARS-CoV-2 infections [41].

To investigate whether SARS-CoV-2 had been introduced into the European wild ruminant population, as occurred in North American white-tailed deer, we serologically tested samples collected in Germany in autumn and winter 2021/22 from various species. In addition, we included the livestock pathogens BTV, SBV and ruminant pestiviruses in the serosurvey.

## Materials and methods

### Wildlife samples

Between September 2021 and January 2022, blood samples of fallow deer (n=26), red deer (n=188), roe deer (n=254), mouflons (n=7) and European bison (Wisent) (n=1) were collected post-mortem in five German federal states by local hunters (Table 1). Further 17 wild ruminant samples were analysed, but the species was not indicated in the letter accompanying the samples. The federal states comprised Bavaria, Hesse (3 hunting districts), Mecklenburg-Western Pomerania, North Rhine-Westphalia (7 hunting districts) and Rhineland-Palatinate (6 hunting districts). The wisent, two of the mouflons, two red deer and 12 fallow deer were kept in wildlife parks, the other ruminants were free-ranging. Control samples collected in the same regions prior to the human SARS-CoV-2 pandemic, i.e. before 2020, were not available.

**Table 1:**
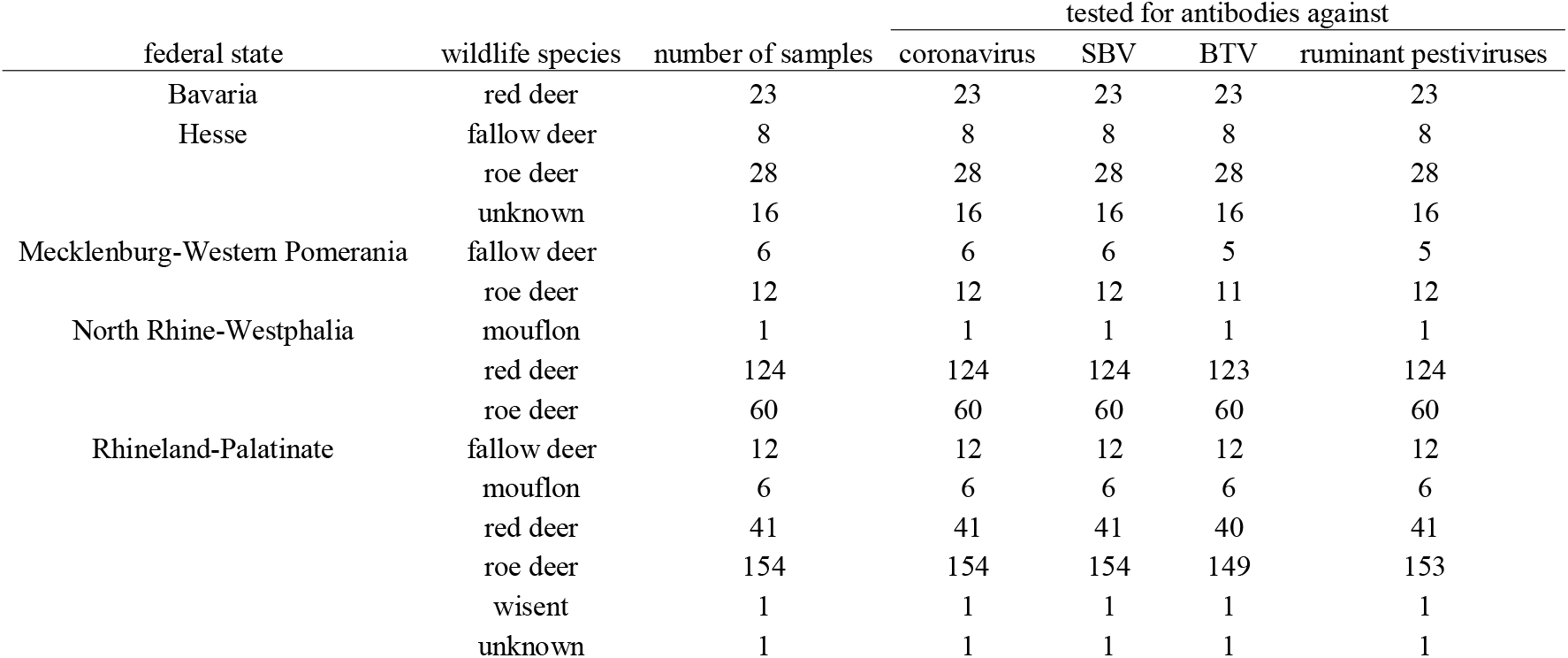
Origin and number of ruminant samples collected in the 2021/22 hunting season in Germany.

In addition, 307 wild ruminant samples collected between 2017 and 2020 at three military training areas of the German Federal Armed Forces were included. In training area A, red deer and roe deer samples were taken in 2017 and in 2019. In training area B, samples were continuously collected from 2017 to 2020 and from training area C, samples from 2017, 2018 and 2019 were available (Table 2).

**Table 2.**
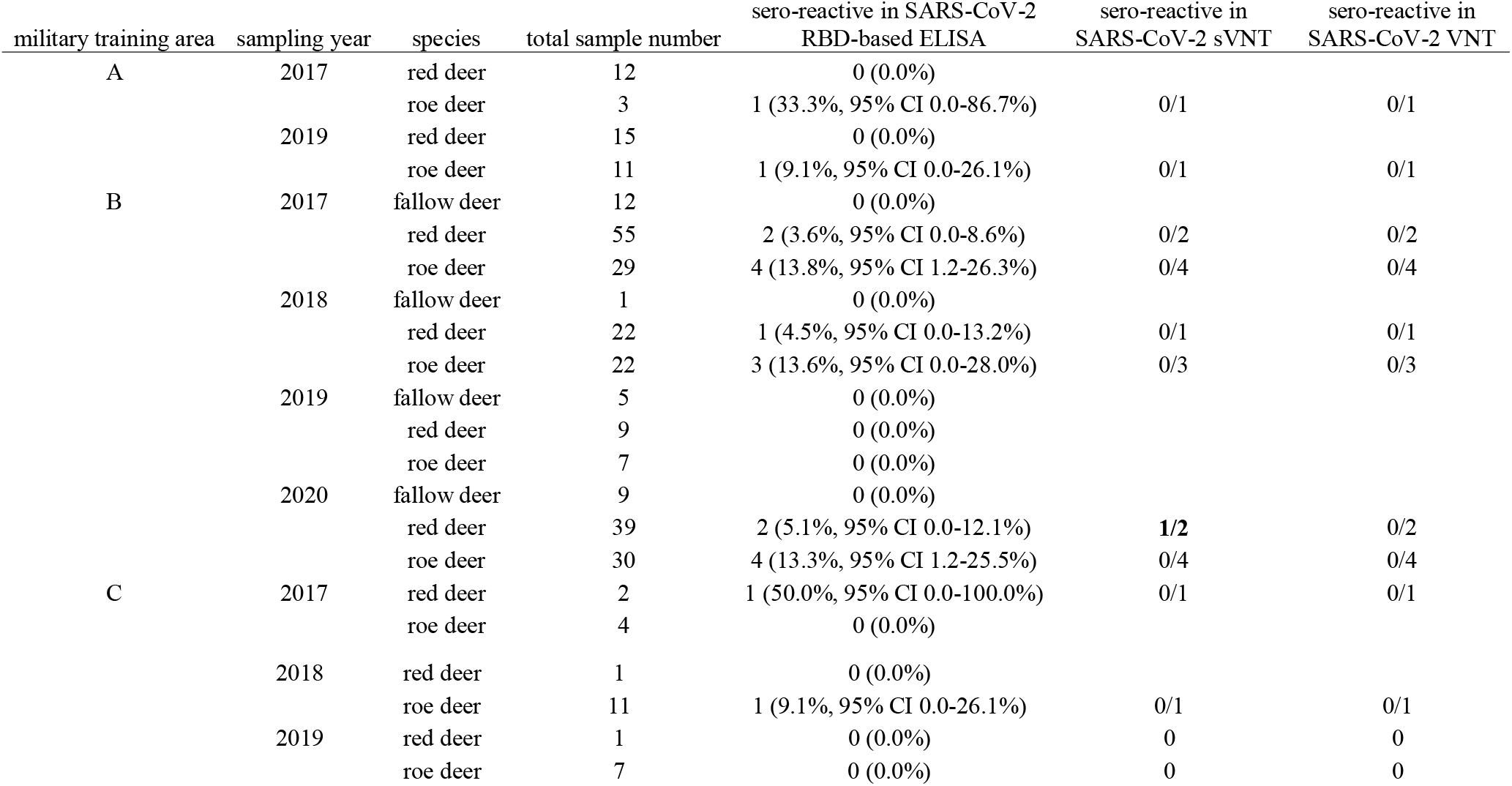
Results of the SARS-CoV-2 serological assays for wild ruminant samples collected at military training areas of the German Federal Armed Forces before and during the COVID-19 pandemic. Sera that reacted positive in the RBD-based ELISA were subsequently analysed by a virus neutralisation test (VNT) and by a surrogate virus neutralisation test (sVNT). CI – confidence interval

### Serological test systems

All samples collected in the 2021/22 hunting season were tested for the presence of anti-SBV antibodies by a glycoprotein Gc-based ELISA performed as described previously [42]. For the detection of antibodies against BTV and ruminant pestiviruses the VP7-based ID Screen Bluetongue Competition ELISA and the ID Screen BVDV p80 Ab competition test, respectively, were applied (both Innovative Diagnostics, Grabels, France). The latter allows for the detection of anti-BVDV antibodies as well as for antibodies against the ovine BDV. Both tests were performed as prescribed by the manufacturer.

The complete sample panel was analysed by a previously described multispecies ELISA based on the receptor-binding domain (RBD) of SARS-CoV-2 [43]. A corrected optical density (OD) of ≥0.3 was evaluated as being positive as determined by prior validation [43]. Samples that reacted positive in the RBD-ELISA were subsequently tested by a virus neutralization test (VNT) against the SARS-CoV-2 strain 2019_nCoV Muc-IMB-1 performed as described previously [44], and by a surrogate VNT (cPass SARS-CoV-2 Surrogate Virus Neutralization Test (sVNT) Kit, GenScript, the Netherlands).

For the detection of antibodies directed against SARS-CoV-1, an ELISA system was established in line with the SARS-CoV-2 test. For expression of the recombinant protein, the RBD-SD1 domain (aa 306 – 577) of the SARS coronavirus strain Tor2 (NC_004718.3) was ordered as a synthetic DNA string fragment (GeneArt synthesis; Thermo Fisher Scientific, Darmstadt, Germany) and cloned into the pEXPR103 expression vector (IBA Lifesciences, Göttingen, Germany) in-frame with a C-terminal Strep-tag. Expi293 cells were grown in suspension in Expi293 expression medium (Thermo Fisher Scientific) at 37 °C, 8 % CO_2_ and 125 rpm. For transfection the ExpiFectamine293 transfection kit (Thermo Fisher Scientific) was used according to the manufacturer’s instructions. Cell culture supernatant was harvested 6 days after transfection and purified using Strep-Tactin XT Superflow high capacity resin (IBA Lifesciences) following the manufacturer’s instructions. The ELISA procedure was exactly as described for the SARS-CoV-2 RBD-ELISA [43]. The adsorbance value was calculated by subtracting the OD value measured at a wavelength of 450 nm on the uncoated well from the value obtained from the protein-coated well for the respective sample.

## Results

### Serology of vector-borne and transboundary livestock diseases

Antibodies against the *Culicoides*-transmitted SBV were detected in 67 of the 493 wildlife samples collected in 2021/22 (13.6 %, 95% confidence interval (CI): 10.6 % - 16.6 %). With exception of the European bison (Wisent), individual animals of every species sampled in the western part of Germany were affected (Figure 1). In contrast, all but one of the analysed samples tested negative for antibodies against the likewise *Culicoides*-transmitted BTV (1/484 positive; 0.2 %, 95% CI: 0 % - 0.6 %). The remaining 9 samples (1 fallow deer, 2 red deer, 6 roe deer) could not be tested due to a lack of sample material. The ELISA-positive sample originated from a fallow deer of unknown age that was hunted in the federal state Hesse (Figure 1).

**Figure 1.**
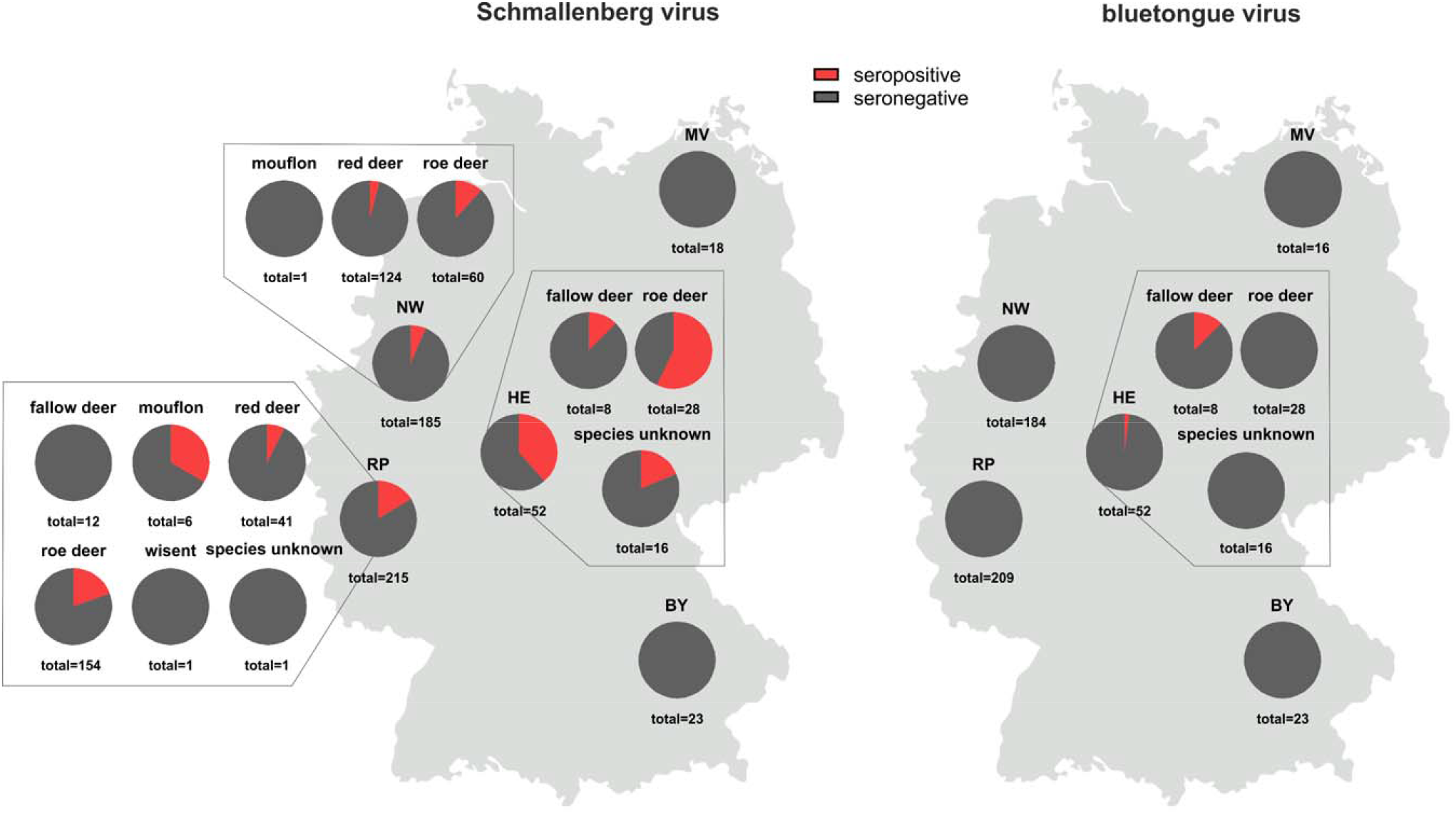
Proportion of wild ruminant samples that tested positive (red) for antibodies against the *Culicoides*-transmitted viruses Schmallenberg virus (left) and bluetongue virus (right). BY – Bavaria, HE – Hesse, MV – Mecklenburg-Western Pomerania, NW – North Rhine-Westphalia, RP – Rhineland-Palatinate

All but two samples of the panel collected in 2021/22 (1 fallow deer and 1 roe deer), which could not be tested because of insufficient sample volume, were additionally analysed for antibodies against ruminant pestiviruses and all of them gave negative results (0/491 positive; 0 %).

### Evidence for the presence of a coronavirus of the Sarbecovirus subgenus

The 493 samples collected in the 2021/22 hunting season were tested by a SARS-CoV-2 RBD-based ELISA and 25 reacted positive (25/493; 5.1 %, 95% CI: 3.1 % - 7.0 %). Two of the sero-reactive samples originated from red deer (2/188), 22 from roe deer (22/254) and for one sample the species was not indicated (1/17) (Figure 2). However, in the surrogate VNT, which was used as confirmatory test, only one red deer serum collected in North Rhine-Westphalia tested positive (31 % inhibition, cut-off for positivity at ≥ 30 % inhibition). In the second confirmatory test, i.e. the VNT, using replicating SARS-CoV-2, none of the sera tested positive.

**Figure 2.**
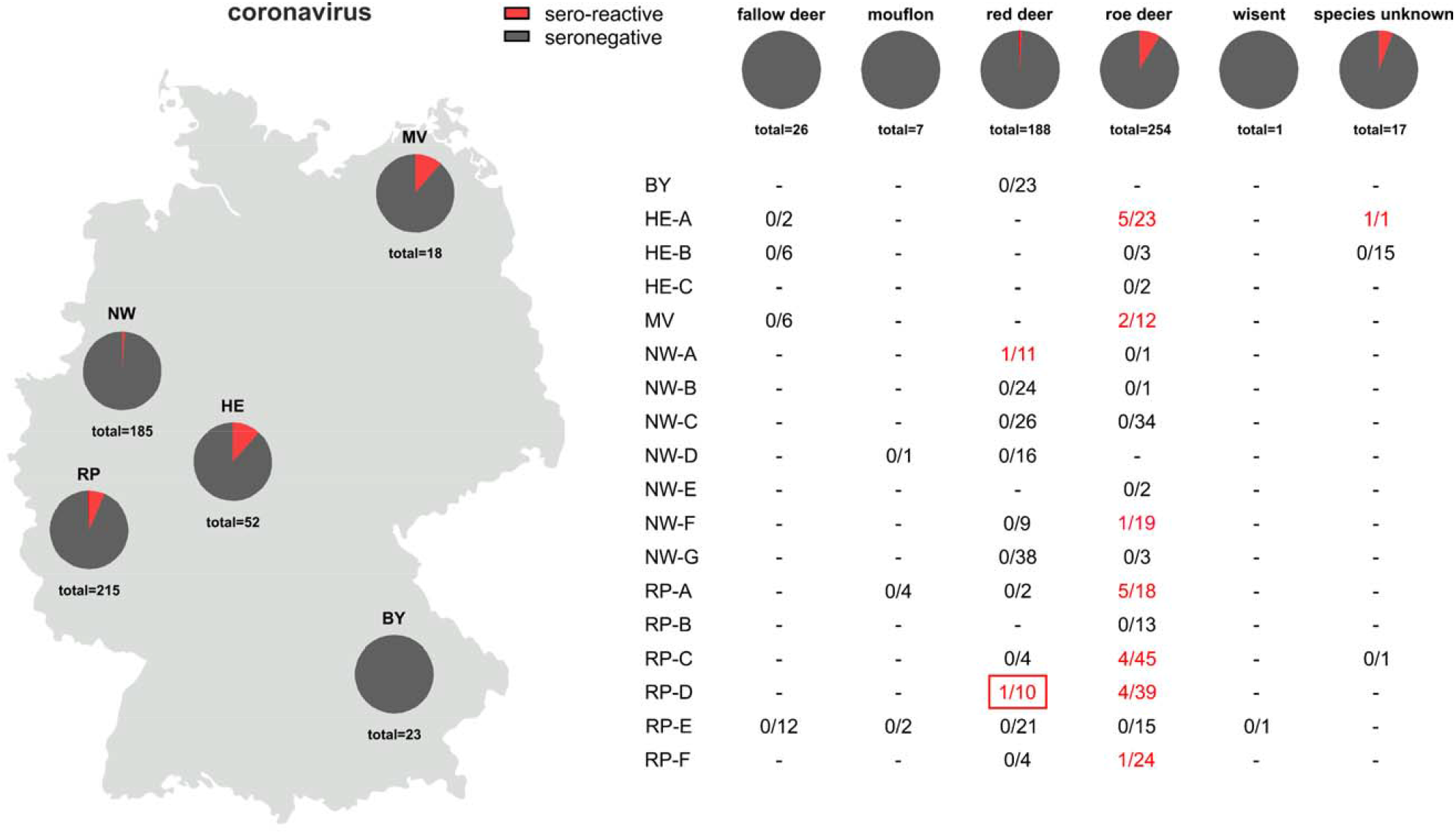
Proportion of samples per German federal state (left) and per wild ruminant species (right) that reacted positive (red) in an RBD-based SARS-CoV-2 antibody ELISA. In the lower left panel, the number of sero-reactive results/number of analysed samples is given separately for each federal state and, if known, the hunting district from which the samples originated. The serum that additionally tested positive in a surrogate virus neutralisation test is framed in red. BY – Bavaria, HE – Hesse, MV – Mecklenburg-Western Pomerania, NW – North Rhine-Westphalia, RP – Rhineland-Palatinate

The observed RBD-reactivity was further examined by using wildlife samples collected at the military training areas before and during the SARS-CoV-2 pandemic and reactive sample were found at every sampling time point (Table 2). A total of 20 out of 307 sera tested positive in the RBD-based ELISA (6.5 %, 95% CI: 3.8 % - 9.3 %) and again, only one of these results could be confirmed by the surrogate VNT (44 % inhibition) and none by the cell-culture based VNT (Table 2). To further investigate whether the reactivity against the SARS-CoV-2-RBD could be induced by antibodies against a hitherto unknown deer coronavirus of the *Sarbecovirus* subgenus, the reactive samples of the panel collected between 2017 and 2020 at the military training areas were additionally tested by an indirect ELISA against the RBD of SARS-CoV-1. 18 of 20 sera scored positive (corrected OD ≥0.3) (Figure 3). As SARS-CoV-2-specific controls, a serum sample obtained from a cattle after experimental SARS-CoV-2 infection [35] and a bovine serum taken after natural infection [38] were analysed and both samples tested positive in the SARS-CoV-2 RBD ELISA but, in contrast to the wildlife samples, negative against SARS-CoV-1 (Figure 3).

**Figure 3.**
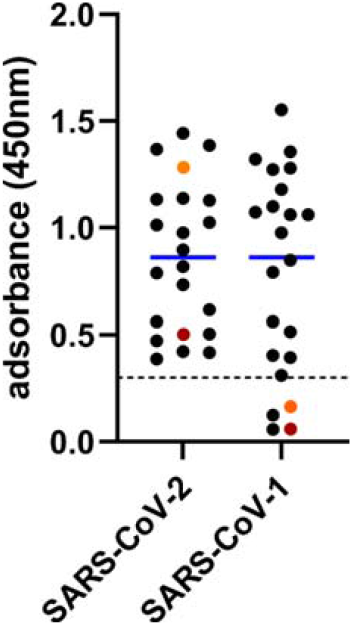
Reactivity of wild ruminant samples collected between 2017 and 2020 towards the receptor-binding domains of the coronaviruses SARS-CoV-2 and SARS-CoV-1 as measured by indirect multispecies ELISAs. The cut-off for positivity (≥0.3) is marked by a dashed horizontal line and the mean values are shown in blue. A cattle sample taken 20 days after experimental SARS-Cov-2 infection (dark red) and a cattle sample taken subsequent to natural SARS-CoV-2 infection (orange) were used as virus-specific controls.

## Discussion

The human SARS-CoV-2 pandemic with hundreds of millions of infected individuals worldwide is presently driven by direct human-to-human virus transmission via aerosolized particles. However, when the virus was introduced into mink farms resulting in local epidemics in these highly susceptible animal species, evidence for mink-to-human spillback infections have been reported [45], resulting in concerns about viral maintenance in animals. Aside from these mink-to-human and the likely initial animal-to-human transmissions, there are currently no further reports of zooanthroponotic SARS-CoV-2 transmissions. Nevertheless, the frequent virus detections in the white-tailed deer population and the presumed intra-species transmission [29–32,40] could pose a risk for infections of humans with close contact to these animals, like e.g. hunters, and also for the establishment of a novel SARS-CoV-2 animal reservoir allowing the virus to continuously mutate with the potential of development of novel variants of concern also for humans. Here, we investigated by serological methods whether SARS-CoV-2 has been introduced into European wild ruminants as has been into the North American white-tailed deer population.

Serological investigations of wildlife animals represent a cornerstone of disease surveillance [46], especially when virus shedding is only transient and the timeframe unknown, since antibodies remain detectable for longer periods. However, when conducting sero-epidemiological studies, potential cross-reactivity between closely related viruses needs to be considered, in the present study in particular between SARS-CoV-2 and other coronaviruses. Here, we found an anti-RBD-reactivity in pre- and post-pandemic wild ruminant sera that could not be confirmed by the highly specific VNT and only in two cases by a surrogate VNT, which is likewise based on the RBD, more precisely on the interaction between SARS-CoV-2’s RBD and the human host cell receptor protein ACE2. In roe deer, which most frequently showed an anti-RBD reactivity in this study, the coronavirus prevalence is largely unknown, but in some other cervid species, bovine-like coronaviruses were found [47–50]. During the initial validation of the RBD-based ELISA used in this study and during an experimental SARS-CoV-2 infection study in cattle, it could be shown that the ELISA does not exhibit cross-reactivity with bovine coronavirus (BCoV) [35,43]. Hence, there is most likely at least one hitherto unknown coronavirus present in the deer population that is more closely related to viruses of the *Sarbecovirus* subgenus than BCoV, leading to the generation of antibodies cross-reactive with both, SARS-CoV-1 and SARS-CoV-2. Interestingly, such an RBD-reactivity that could not be explained by circulation of known coronaviruses was also observed in sera of domestic and peridomestic animals collected in the United States prior to the current SARS-CoV-2 pandemic [51]. In addition, SARS-CoV-2 reactive antibodies have been rarely found in pre-pandemic human sera, but these antibodies target the S2 subunit of the spike protein [52]. The S1 subunit, which represents the second subunit of which the spike protein is composed, contains the RBD that is responsible for virus interaction with the host cell receptor protein ACE2 reflecting virus entry and host specificity [53]. S1 including the RBD is much more divergent than S2 between SARS-CoV-2 and other circulating coronaviruses of humans and animals, making it a highly specific target for serological test systems [54]. The RBDs of SARS-CoV-1 and SARS-CoV-2, however, exhibit major similarities in protein structure as well as sequence and cross-reactions occur [55]. Hence, the cross-reacting agent found in German wild ruminants is most likely a coronavirus of the *Sarbecovirus* subgenus closely related to both, SARS-CoV-1 and SARS-CoV-2. Nevertheless, for final classification of the cross-reacting coronavirus, the identification of the virus in question by PCR methods and/or sequence analysis of tissue samples is required.

In contrast to SARS-CoV-2, which was discovered only about two years ago, the host range of the livestock diseases that were included in this study is largely known and wild ruminants are part of it. Regarding pestiviruses, PI animals have been previously identified in wild ruminant species [26,27], but in this study we did not detect any seropositive animal, making BVDV or BDV infections and in particular the presence of PI animals highly unlikely in the population surveyed. Therefore, it is very improbable that wildlife species form a significant virus reservoir and could be a source of infection for domestic cattle, which is nearly free from the infection as a result of the mandatory control program (0.005 % PI prevalence among all newborn calves in 2020 [56]).

For the arbovirus SBV a different situation was observed, since about 13.6 % of the sampled animals tested seropositive, which mirrors the situation in domestic ruminants. In the absence of control measures, SBV established an enzootic status after its emergence in Central Europe a decade ago [12,14,15]. In domestic animals, a wavelike pattern of circulation with increased case numbers every two to three years has been observed [14,15]. Unfortunately, the age of the wild ruminants sampled for the present study is known for only a subset of animals, which hampers a more detailed analysis of the year of infection. Nevertheless, the dataset allows a comparison between the infection rates with the enzootic SBV and BTV, for which sporadic cases have been reported in domestic ruminants since December 2018 with the last case reported in February 2021 [18]. Both viruses have their mammalian hosts and the insect vector species responsible for virus transmission in common [9,10], and, hence, they share major factors for virus circulation in a given area. Whereas anti-SBV antibodies were detected rather frequently in samples collected in the 2021/22 hunting season, only one animal tested positive in the BTV ELISA. The age of this animal is not known, but given the epidemiological situation in domestic ruminants [18] and the lack of further positive results, the seropositive fallow deer was most likely older than one year and was infected in one of the previous vector seasons. The differences in the seroprevalences of antibodies against SBV and BTV in the wild ruminant population and the agreement with the respective situation in domestic animals confirm previous observations that both, wildlife and farmed animals, are part of the transmission cycle, but that wild ruminants do not play a significant role in the maintenance of BTV in a given area [57,58]. For SBV, similar spatio-temporal distribution patterns were previously found in wild ruminants following outbreaks in domestic animals [6,8,59]. BTV or BTV-specific antibodies were detected in wild ruminants only in regions where the virus is also present in domestic animals [57]. Despite the presence of competent insect vectors, BTV serotypes 1 and 8 never spread in red deer beyond the domestic outbreak range in France [58]. Here, the single seropositive animal was hunted in a federal state within the restricted zone, while anti-SBV antibodies are widespread, highlighting the suitability of serological surveys in wildlife animals to demonstrate presence or absence of multi-host diseases in a given area [46].

## Acknowledgements

We thank Bianka Hillmann, Mareen Lange and Antje Gromtzik for excellent technical assistance and the hunters for providing the wildlife samples.

## Ethical Statement

Blood samples were collected by local hunters according to the appropriate German legislation. No ethical/welfare authority approval was required as all samples were collected post-mortem by the hunters.

## Data availability statement

The data that support the findings of this study are available from the corresponding authors, KW and MB, upon reasonable request.

